# Ataxia-linked SLC1A3 mutations alter EAAT1 chloride channel activity and glial regulation of CNS function

**DOI:** 10.1101/2021.10.05.463128

**Authors:** Qianyi Wu, Azman Akhter, Shashank Pant, Eunjoo Cho, Jin Xin Zhu, Alastair Garner, Tomoko Ohyama, Emad Tajkhorshid, Donald J. van Meyel, Renae M. Ryan

**Author notes:** These authors share joint first authorship (Qianyi Wu, Azman Akhter).

## Abstract

Glutamate is the predominant excitatory neurotransmitter in the mammalian central nervous system (CNS). Excitatory Amino Acid Transporters (EAATs) regulate extracellular glutamate by transporting it into cells, mostly glia, to terminate neurotransmission and to avoid neurotoxicity. EAATs are also chloride (Cl^−^) channels, but the physiological role of Cl^−^ conductance through EAATs is poorly understood. Mutations of human EAAT1 (hEAAT1) have been identified in patients with episodic ataxia type 6 (EA6). One mutation showed increased Cl^−^ channel activity and decreased glutamate transport, but the relative contributions of each function of hEAAT1 to mechanisms underlying the pathology of EA6 remain unclear. Here we investigated the effects of five additional EA6-related mutations on hEAAT1 function in *Xenopus laevis* oocytes, and on CNS function in a *Drosophila melanogaster* model of locomotor behavior. Our results indicate that mutations with decreased hEAAT1 Cl^−^ channel activity and functional glutamate transport can also contribute to the pathology of EA6, highlighting the importance of Cl^−^ homeostasis in glial cells for proper CNS function. We also identified a novel mechanism involving an ectopic sodium (Na^+^) leak conductance in glial cells. Together, these results strongly support the idea that EA6 is primarily an ion channelopathy of CNS glia.

## Introduction

Episodic ataxias (EA) are a group of rare neurological disorders characterized by progressive, severe and recurrent episodes of ataxia, migraine, discoordination, and imbalance (1). Nine types of EA (EA1-9) have been identified (2–11), the most common of which are EA1 and EA2. EA1 patients have mutations in *KCNA1*, a gene that encodes potassium channel Kv1.1 (3). EA2 is characterized by mutations in *CACNA1A* which encodes the calcium channel Cav2 (6), and EA5 results from mutations in *CACNB4* (12). While EA1, EA2, and EA5 are directly related to mutations in ion channels, EA8 and EA9 are associated with ion channel dysfunction, and no candidate genes have been reported for EA3, EA4 and EA7 [for a review see Maksemous, et al. (10)]. EA6 patients have mutations in *SLC1A3*, the gene which encodes the human <u>E</u>xcitatory <u>A</u>mino <u>A</u>cid <u>T</u>ransporter 1 (hEAAT1). The etiology of EA6 likely traces to Bergmann glia, which are astrocytes in the cerebellum that express EAAT1 at high levels and envelop the dendritic arbors of Purkinje neurons (13).

Glutamate is the predominant excitatory neurotransmitter in the central nervous system (CNS) (14–16), with concentrations estimated to be as low as 25 nM at rest (17) and rising to the millimolar range upon activation of glutamatergic neurons (18, 19). These elevated levels are rapidly cleared by the EAATs that transport glutamate into nearby glial cells (20–24). Tight regulation of extracellular glutamate levels is important to maintain dynamic signaling between neurons and to avoid neural toxicity (20, 21, 25–27).

The transport of glutamate via EAATs is coupled to the co-transport of three sodium (Na^+^) ions, a proton (H^+^), and the counter-transport of a potassium (K^+^) ion (28, 29). The binding of Na^+^ and glutamate to the EAATs also activates a thermodynamically uncoupled chloride (Cl^−^) conductance (26, 30, 31) (Fig. 1A). The physiological role of this EAAT-dependent Cl^−^ flux has been implicated in charge neutralization, ionic homeostasis and regulation of presynaptic glutamate release but is yet to be fully elucidated (32–35).

**Figure 1.**
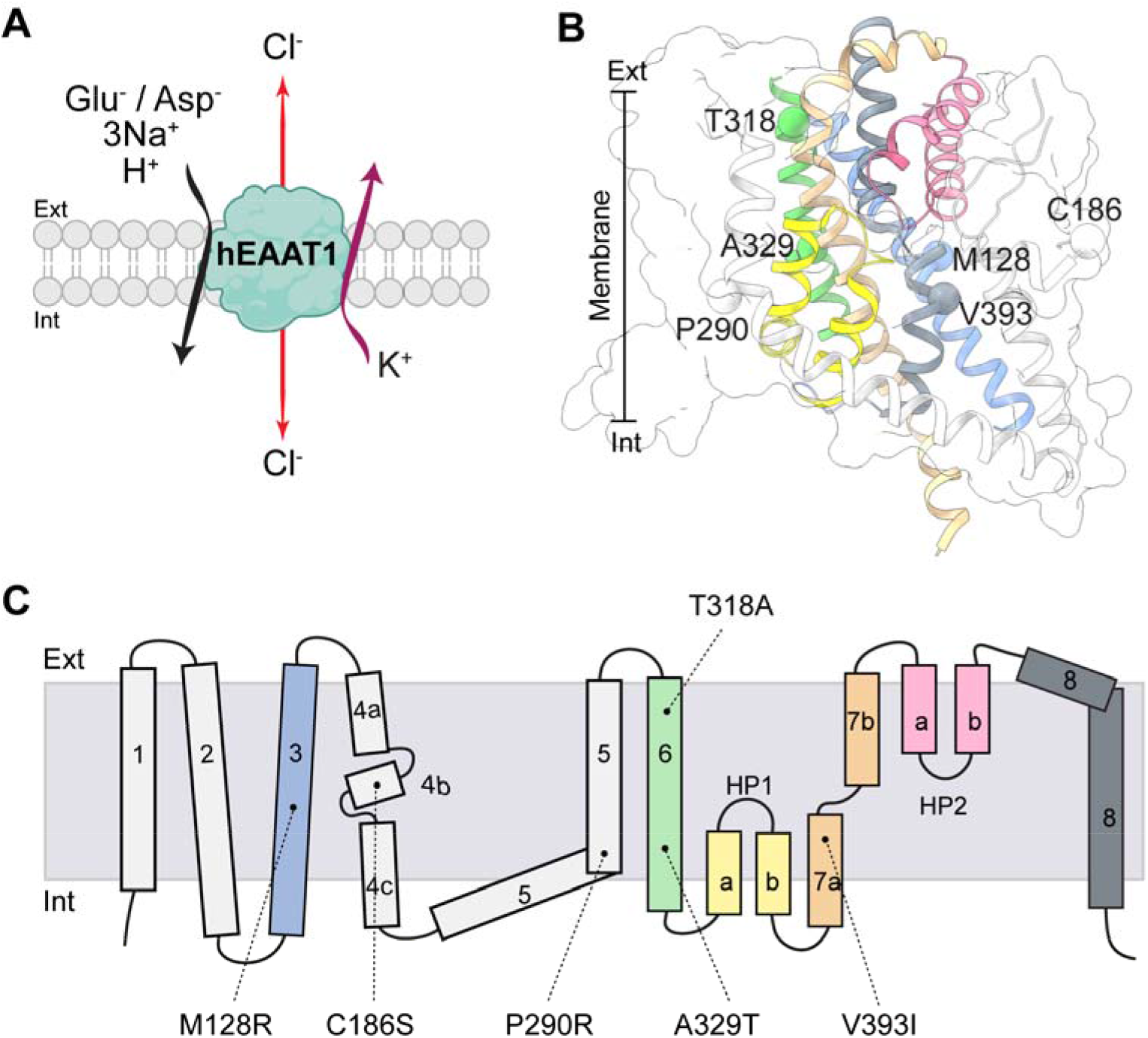
Stoichiometry of transport and locations of EA6 mutations in the hEAAT1 structure. (**A**) The transport of each substrate molecule (glutamate or aspartate) is coupled to the co-transport of three sodium ions (Na^+^) and one proton (H^+^), and the counter-transport of a potassium ion (K^+^); the transport process also activates a thermodynamically uncoupled chloride channel. **(B, C)** Locations of EA6-related mutations in the structure of hEAAT1 (PDB5LLU) (B) and on a topology schematic (C) are shown. Cartoon representation of transmembrane domains (TMDs) TM1 and TM2 from the scaffold domain are omitted for clarity, whilst the whole scaffold domain is shown in transparent white surface representation. TMDs in (B) are coloured as per (C). Figure was made using ChimeraX (Pettersen *et al.*, 2021).

Several mutations in hEAAT1 have been reported in EA6 patients with variability in clinical features such as the age of disease onset, ictal symptoms and duration of attacks (7, 12, 36–38). EA6 was first identified in a child with early onset episodes of ataxia, hemiplegia, seizures, migraine, and epilepsy, in whom a single point mutation, c.869C>G (P290R), was identified in *SLC1A3* (7). P290R resulted in reduced expression of hEAAT1 at the cell surface, reduced rate of glutamate transport, and increased Cl^−^ channel activity (7, 39, 40). Several other EA6-related mutations have since been identified in hEAAT1, including M128R, C186S, T318A, A329T, and V393I, which were proposed to be disease-causing in genomic studies (12, 36–38). While the binding sites for substrate and co-transported Na^+^ ions are well understood in the SLC1A family (41–44), as is the location of the Cl^−^ channel (31, 45, 46), the EA6-related mutations are located in distinct regions of the hEAAT1 protein (Fig. 1B,C), and so the link between these hEAAT1 mutations and the pathogenesis of EA6 is unclear.

*Drosophila melanogaster* provides robust rescue assay of larval locomotion to study the functional impact of EA6-related mutations of hEAAT1 *in vivo*. The larval CNS controls crawling behavior using many of the same neurotransmitters as humans, including glutamate, and is comprised of two brain lobes and a ventral nerve cord (VNC) in which there are roughly 10,000 neurons and glial cells, including astrocytes. *Drosophila* astrocytes, like their counterparts in humans, express high levels of EAAT1 (dEAAT1) (47), have ramified arbors that infiltrate the synaptic neuropil (48, 49), and play an important role in modulating the functions of synapses within CNS circuits by regulating ion and neurotransmitter homeostasis. *Drosophila* larvae normally move via peristaltic crawling interrupted occasionally by brief pauses and turning. dEAAT1 is essential for larval crawling (49), as *dEAAT1*-null larvae rarely make these peristaltic contractions. This defect can be rescued by expressing the human ortholog hEAAT1 in larval CNS astrocytes using the GAL4-UAS-system (48, 49). This rescue model revealed that EAAT1 function is conserved between *Drosophila* and humans. However, hEAAT1 bearing the P290R mutation could not rescue the crawling defects of *dEAAT1*-null larvae, indicating this particular mutation renders hEAAT1 nonfunctional *in vivo* (48). The P290R mutation had both reduced glutamate transport (7) and a large increase in Cl^−^ channel activity (39), and so the relative contribution of each function to EA6 pathology was unclear. However, co-expression of other Cl^−^ transport proteins in this Drosophila model provided strong evidence for the importance of the Cl^−^ channel activity (48), and subsequently a mouse model for the P290R mutation demonstrated increased glutamate-activated Cl^−^ efflux from Bergmann glia that triggers apoptosis in the cerebellar cortex (50).

In this study, we investigated the functional impact of five additional hEAAT1 mutations related to EA6. We found hEAAT1 Cl^−^ channel activity is essential for CNS function *in vivo* and we linked altered Cl^−^ channel activity of the EA6-related mutations with deficits in *Drosophila* motor behavior. Using a combination of functional analysis and molecular dynamics simulations we found that one severe mutation (M128R) also altered protein/membrane interactions and introduced an ectopic sodium (Na^+^) leak conductance. Together, these results strongly support the idea that EA6 is primarily an ion channelopathy of CNS glia involving disrupted homeostasis of Cl^−^, though unusual Na^+^ flux or glutamate transport may also contribute in some instances.

## Results

### Expressing hEAAT1 and EA6-related mutant transporters in Drosophila larvae

With the GAL4-UAS system and *alrm-Gal4* for selective expression of hEAAT1 in larval CNS astrocytes, we used infrared video tracking to examine the effects of five EA6-related mutations (M128R, C186S, T318A, A329T, and V393I) on the ability of hEAAT1 to rescue the larval crawling defects of *dEAAT1*-null animals. Tracking videos for control animals and those bearing each of the EA6-related mutations revealed a range of differences in their ability to rescue *dEAAT1*-nulls (Supplementary Movies 1-7). This range is reflected in representative reconstructions of larval crawling paths and velocities (9 animals/genotype, Fig. 2A). We quantified the performance of animals in each genotype during 60s of free exploration and found that, like hEAAT1, the C186S and T318A mutations could rescue key features of larval crawling behavior, including the mean speed (Fig. 2B), the total path length achieved in 60s (Fig. 2C), and the beeline distance from origin reached in 60s (Fig. 2D). In contrast, the M128R and V393I mutations could not rescue any of these features, and the A329T rescued them only partially (Fig. 2A–D).

**Figure 2.**
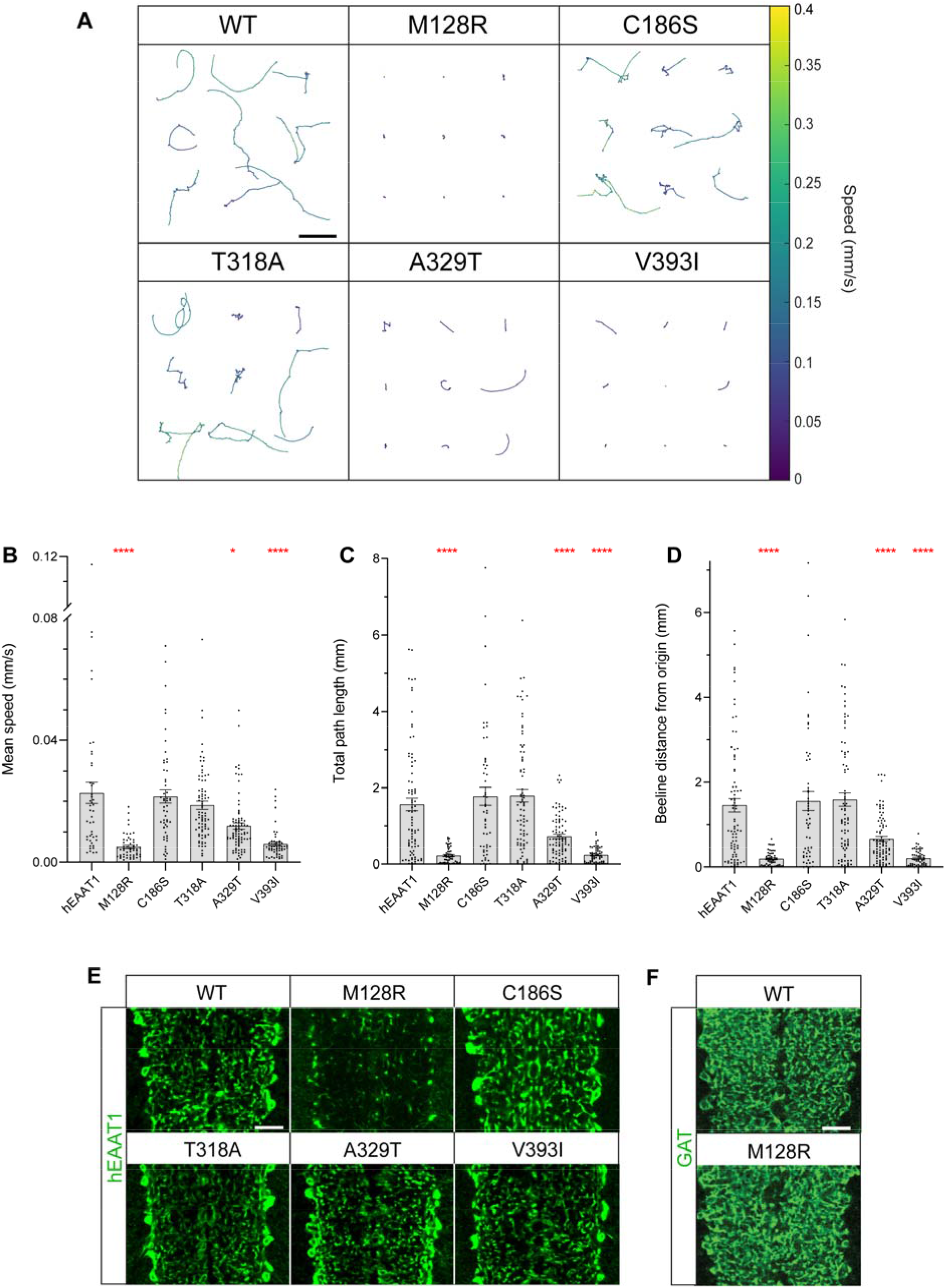
EA6-related mutations M128R, A329T and V393I fail to rescue *Drosophila* crawling. **(A**) Representative trajectories of crawling paths of L1 larvae captured by infrared tracking for 180 s and analysis with MWT software. A colour heatmap indicates the average speed over a moving bin of 0.5 s. Paths of nine larvae are shown for each genotype where *dEAAT1*-null animals were rescued by hEAAT1-WT or one of five EA6-related mutations (M128R, C186S, T318A, A329T or V393I). Scale bar (5 mm) in hEAAT1 panel applies to other panels also. **(B-D)** Quantification in bar graphs (mean ± standard error (SEM)) of crawling parameters achieved by larvae over 60 s of continuous tracking, including mean speed (B), total path length (C), and the beeline distance between the origin at *t* = 0 and the termination point (D). One-way ANOVA tests (Brown-Forsythe) were performed for mean speed F(5,130) = 18.32, p < 0.0001, for total path length F(5,197.6) = 27.16, p < 0.0001, and for beeline distance F(5,199.5) = 24.04, p < 0.0001. **(E, F)** Astrocyte-specific expression (with alrm-Gal4) of hEAAT1 and EA6-related mutations. Representative high-power images of infiltrative astrocyte processes within the ventral nerve cord of a dissected larva for each genotype, labelled with immunohistochemistry for hEAAT1 (E) or the plasma membrane-associated GABA transporter (Gat, F). Each panel represents a single optical confocal section from the middle of the dorsal-ventral axis of the neuropil. Scale bars =10 μm. Like the control where *dEaat1*-null larvae are rescued with hEAAT1, all the EA6-related mutations of hEAAT1 except M128R are well expressed and addressed to astrocyte processes within CNS neuropil. Anti-Gat staining (F) reveals that astrocytes rescued with M128R infiltrate the neuropil normally.

Except for M128R, each of the mutant transporters was strongly expressed in astrocytes and their ramified processes throughout the neuropil, as observed with immunohistochemistry using an antibody specific to hEAAT1 (Fig. 2E). M128R was not well expressed there, and so we used immunohistochemistry for the GABA transporter (Gat), which is expressed on the surface of astrocytes, to confirm that the reduced expression of M128R was not due to a failure of astrocytes to infiltrate neuropil in this genotype (Fig. 2F). Instead, it is likely that the M128R mutation limits the expression of hEAAT1 or its distribution to astrocyte processes.

### Expression and functional analysis of EA6-related hEAAT1 mutants in oocytes

The larval locomotion assays indicated that M128R, A329T and V393I disrupted hEAAT1 function, while C186S and T318A did not. To investigate the functional impact of these EA6 mutations, and the previously characterized P290R (7, 39, 48), in an isolated system, they were expressed in *Xenopus laevis* oocytes. Surface expression was monitored by attaching green fluorescent protein (GFP, Supplementary Fig. S1A), which does not affect the function of hEAAT1 (Supplementary Fig. S1C). While most of the EA6-mutants displayed comparable levels of surface expression to hEAAT1 the expression of the P290R mutant was significantly reduced (27.4 ± 4.0 % of hEAAT1, Supplementary Fig. S1B), agreeing with previous reports in HEK and COS7 cells (7, 39). The surface expression of M128R was also significantly reduced (25.9 ± 2.0 % of hEAAT1, Supplementary Fig. S1B), however it was expressed at similar levels as P290R, for which substrate transport can be measured (Figure 3B–D).

**Figure 3.**
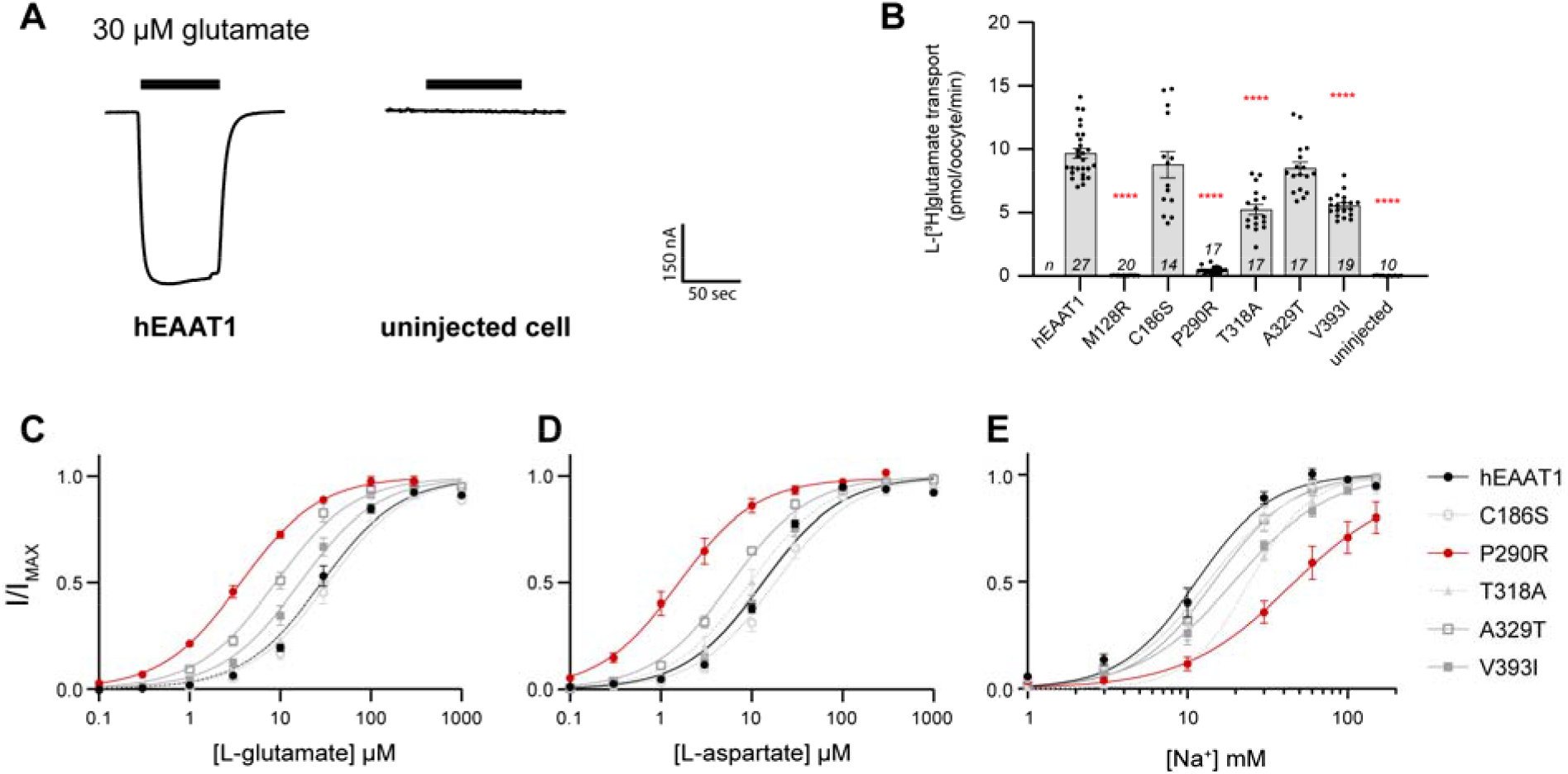
Dose-response relationships for substrates and rate of l-[^3^H]glutamate transport via hEAAT1-WT and EA6-related mutant transporters. (**A**)The hEAAT1 glutamate transport process is associated with the net influx of two positive charges, therefore the application of 30 μM L-glutamate results in an inward current in 96 mM NaCl buffer when the membrane potential is clamped at −60 mV, which is absent from uninjected (control) cells. (**B**) Glutamate uptake measured after incubation of oocytes expressing hEAAT1, the various EA6-mutant transporters and uninjected (control) oocytes with radiolabelled L-[^3^H]glutamate for 10 minutes. One-way ANOVA tests (Brown-Forsythe) were performed F(7,35.81) = 90.7, p < 0.0001. (**C**-**E**) Dose-dependent relationships were determined by currents elicited by L-glutamate (C) and L-aspartate (D) at −60 mV. Dose-dependent relationships for Na^+^ were determined with a saturating dose of L-glutamate (300 μM, except for P290R which has a higher affinity for L-glutamate, and for which 100 μM was used) (E). For the exact number of cells (n) used and analysis of electrophysiological properties, see Table 1.

**Table 1.** Electrophysiological properties of wild-type and mutant hEAAT1 transporters. Apparent affinities (*K*_*m*_) of l-glutamate, l-aspartate, and Na^+^ analysed from dose-response curves obtained in Figure 3 C–E and Supplementary Figure 3 B-D are shown. Data represented as mean ± SEM, and the number of cells (n) used is indicated.

| hEAAT1 | $K_m$ ($\mu\text{M}$) | | | | $\text{Na}^+$ | | |
| --- | --- | --- | --- | --- | --- | --- | --- |
| | L-Glu | n | L-Asp | n | $K_m$ (mM) | Hill | n |
| WT | $30.0 \pm 3.4$ | 8 | $13.4 \pm 0.8$ | 5 | $11.0 \pm 1.1$ | $2.4 \pm 0.6$ | 5 |
| M128R | - | - | - | - | - | - | - |
| M128K | $3.9 \pm 0.3$ | 8 | $3.0 \pm 0.7$ | 7 | $33.2 \pm 6.2$ | $1.3 \pm 0.1$ | 10 |
| C186S | $35.8 \pm 6.7$ | 4 | $18.9 \pm 3.0$ | 4 | $13.6 \pm 1.1$ | $2.1 \pm 0.5$ | 5 |
| P290R | $3.7 \pm 0.4$ | 5 | $1.8 \pm 0.5$ | 5 | $62.4 \pm 19.6$ | $1.5 \pm 0.3$ | 7 |
| T318A | $28.1 \pm 1.7$ | 5 | $10.0 \pm 1.5$ | 5 | $24.3 \pm 1.1$ | $2.3 \pm 0.2$ | 5 |
| A329T | $9.4 \pm 1.3$ | 6 | $6.0 \pm 0.4$ | 6 | $15.8 \pm 2.9$ | $2.0 \pm 0.1$ | 5 |
| V393I | $17.5 \pm 2.2$ | 6 | $13.3 \pm 1.7$ | 5 | $19.3 \pm 0.8$ | $1.7 \pm 0.3$ | 5 |

Glutamate transport via the EAATs results in a net influx of two positive charges per transport cycle, which can be measured as an inward current upon glutamate application to oocytes expressing hEAAT1 clamped at −60 mV (Fig. 3A). Apparent affinities (*K*_*m*_) of l-glutamate were determined for hEAAT1 (hEAAT1, *K*_*m*_ = 30.0 ± 3.4 μM) and the EA6-related mutant transporters (Fig. 3C). The mutations C186S (*K*_*m*_ = 35.8 ± 6.7 μM) and T318A (*K*_*m*_ = 28.1 ± 1.7 μM) had marginal effects on the apparent affinity of l-glutamate, while A329T (*K*_*m*_ = 9.4 ± 1.3 μM) and V393I (*K*_*m*_ = 17.5 ± 2.2 μM) both showed an increase in apparent affinity compared to hEAAT1 (Fig. 3 C & Table 1). Mutations P290R and M128R had strong effects, where the l-glutamate apparent affinity for P290R (*K*_*m*_ = 3.7 ± 0.4 μM) was ~10-fold higher than hEAAT1, while the affinity for M128R could not be measured as no substrate-activated currents were detected (Fig. 4B). For each EA6-related mutation, similar effects were observed when the alternative substrate l-aspartate was applied instead of l-glutamate (Fig. 3D & Table 1).

**Figure 4.**
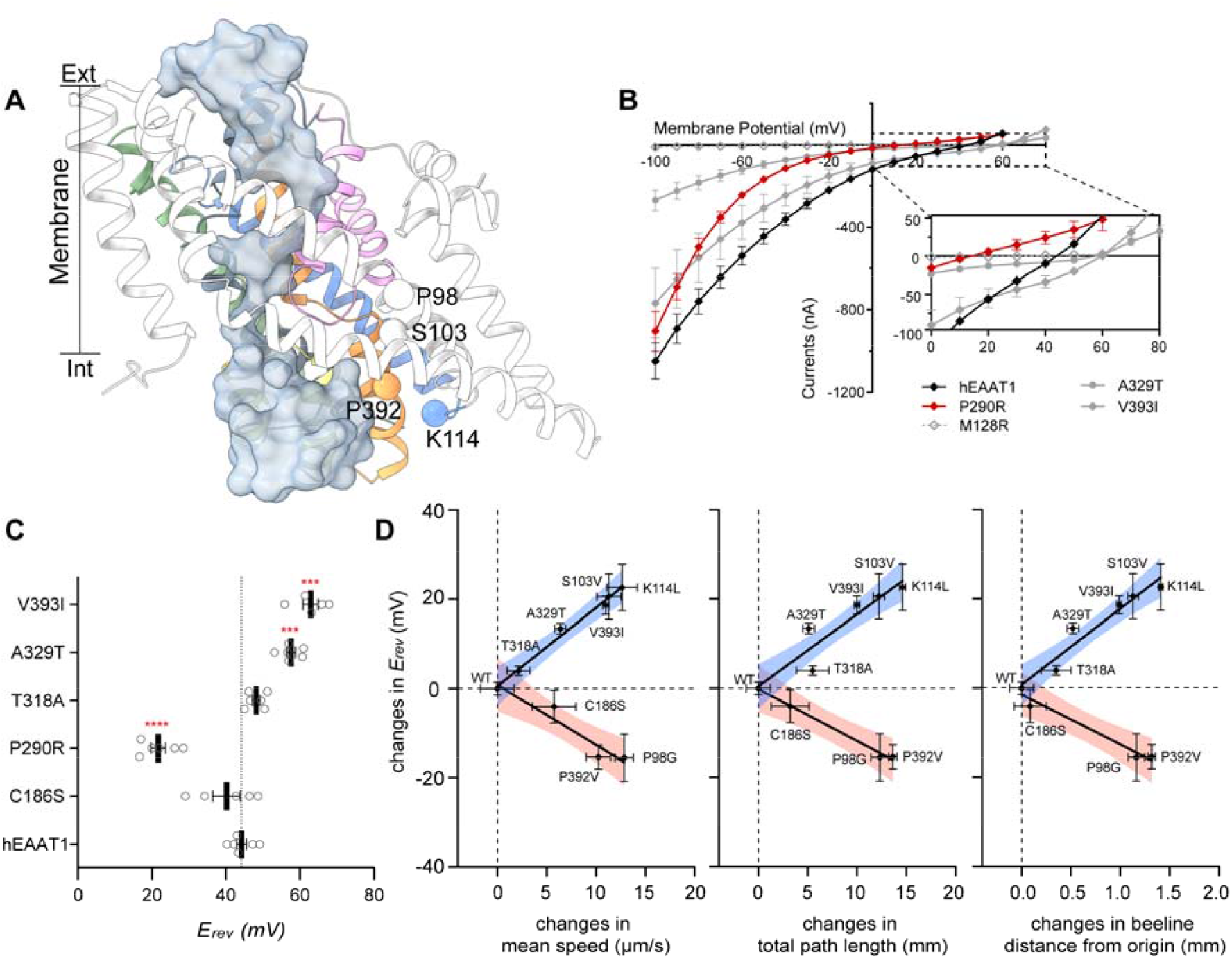
Substrate activated Cl^−^ conductance. (**A**) Positions of residues that, when mutated to the residues indicated, have previously been demonstrated to increase (P98G, P392V) or reduce (S103V, K114L) Cl^−^ conductance in hEAAT1 are highlighted in the structure of glutamate transporter homologue Glt_Ph_ (PDB 6WYK), which was solved in a Cl^−^ conducting state (Cl^−^ pathway indicated as transparent blue density). TMs are coloured as per **Fig. 1 C**. For clarity, hEAAT1 numbering is used for annotation. (**B**) Current-voltage (IV) relationships for hEAAT1 (black diamond), P290R (red diamond), M128R (grey empty diamond), A329T (grey circle) and V393I (grey diamond) were plotted, where reversal potentials (*E*_*rev*_) were measured (**C**). Currents were elicited by 100 μM l-aspartate in 10 mM Cl^−^ buffer containing 100 mM Na^+^ (gluconate was used as Cl^−^ substituent to maintain equal osmolarity), except M128R. *E*_*rev*_ was not determined for M128R due to no detectable l-aspartate activated current, even when 1 mM L-aspartate was applied in the presence of 100 mM NaCl. One-way ANOVA tests (Brown-Forsythe) were performed F(5,13.12) = 50.6, p < 0.0001. (**D**) The change in *E*_*rev*_ of each mutant (compared to hEAAT1), with either increased (*E*_*rev*_ shifted to more negative membrane potentials) or reduced (*E*_*rev*_ shifted to more positive membrane potentials) channel activity, is plotted against the changes in phenotype (including mean speed, total path length, and beeline distance from the origin, **Fig 2** and **5**), and fitted to linear regressions where 95% confidence intervals are indicated as blue or red shadows.

Next, transport of l-[^3^H]glutamate was measured in oocytes expressing hEAAT1 and the EA6-related mutations (Fig. 3B). Compared to hEAAT1, the level of L-[^3^H]glutamate uptake via M128R was not above background levels (uninjected cells), and the P290R mutation was only ~5% of hEAAT1. In contrast, C186S, T318A, A329T and V393I all mediated l-[^3^H]glutamate transport at levels similar to hEAAT1. The P290R mutation is known to affect the ability of Na^+^ to bind hEAAT1 and thus, support the glutamate transport process (39). Therefore, the Na^+^ dependence of glutamate-activated currents for the EA6-related mutations were compared to hEAAT1 (*K*_*m*_ = 11.0 ± 1.1 mM) (Fig. 3E). P290R had the most significant effect on Na^+^ affinity (*K*_*m*_ = 62.4 ± 19.6 mM), about six-fold lower than that of hEAAT1. A small but significant reduction in Na^+^ affinity was observed for T318A (*K*_*m*_ = 24.3 ± 1.1 mM), and V393I (*K*_*m*_ = 19.3 ± 0.8 mM) (Table 1). Taken together, these results indicate that C186S, T318A, A329T and V393I are capable of Na^+^-coupled glutamate transport like that of hEAAT1, while P290R supports low levels of glutamate transport with an increase in affinity for substrate and a concurrent decreased affinity for Na^+^. In contrast, M128R is incapable of any substrate transport or substrate-activated currents, indicating that this mutation severely impacts transport function.

### Substrate-activated uncoupled Cl^−^ conductance

In addition to the currents elicited by Na^+^-coupled glutamate transport, the EAATs have an uncoupled Cl^−^ conductance that is activated upon substrate and Na^+^ binding (14, 30). Previous studies have suggested that the pathology of the P290R mutation is linked to increased Cl^−^ channel activity (39, 48, 50). To explore the contribution of the two components of the transport process, substrate-activated current-voltage relationships (IV) were measured to determine the reversal potential (*E*_*rev*_), which is the membrane potential at which no net flux occurs. The coupled substrate transport conductance for hEAAT1 expressed in oocytes does not reverse at membrane potentials ranging from −100 mV to +60 mV (51), but the Cl^−^ current does reverse at ~-20 mV (*E_Cl-_*) (52), which results in net currents for hEAAT1 with a *E*_*rev*_ = 44.2 ± 1.4 mV (Fig. 4B). To examine whether the uncoupled Cl^−^ channel component of hEAAT1 was affected by the EA6-related mutations, the *E*_*rev*_ of currents elicited by 100 μM L-aspartate was measured (Fig. 4C).

As anticipated, when compared to hEAAT1, the *E*_*rev*_ for P290R (21.7 ± 2.0 mV) was shifted in a negative direction closer to *E_Cl-_* by 22.5 ± 3.4 mV, indicating an increase in Cl^−^ channel function in agreement with previous findings for P290R expressed in HEK293 cells (39). In contrast, the *E*_*rev*_ measured for the mutants A329T (*E*_*rev*_ = 57.6 ± 1.2 mV) and V393I (*E*_*rev*_ = 62.9 ± 2.1 mV) were shifted to more positive membrane potentials, indicating a reduced contribution from Cl^−^, likely due to a decrease in Cl^−^ channel function. Interestingly, in *Drosophila* neither A329T nor V393I could rescue larval crawling behavior (Fig. 2), nor could P290R, as shown previously (48), which increases Cl^−^ channel function. Thus, hEAAT1 function in an intact CNS seems to be compromised by mutations that both decrease and increase Cl^−^ channel function. In support of this link between altered Cl^−^ channel activity and deficits in *Drosophila* motor behavior, the *E*_*rev*_ values of the substrate-activated current of C186S and T318A, which both fully rescued larval crawling, were not significantly different from hEAAT1 (*p* = 0.8714 and 0.2128, respectively) (Fig. 4B and Supplementary Fig. S2).

To explore the idea that too little or too much Cl^−^ flux through hEAAT1 could affect function *in vivo*, and to attempt to distinguish this from the glutamate transport function of hEAAT1, we investigated several well-characterized mutations (Fig. 4A) in our rescue assay for *Drosophila* larval crawling (Fig. 5). None of these mutations have been reported in patients with EA6, but they were used as an independent line of inquiry to examine the importance of the EAAT1 Cl^−^ channel for function *in vivo*. We examined two mutants that have reduced Cl^−^ flux (S103V, K114L) but only minor effects on glutamate transport (45). We found that larvae expressing these mutant transporters were clearly defective in crawling (Fig 5A, and Supplementary Movies 8-11), and in all parameters tested (Fig. 5B–D), even though each of the mutant transporters was strongly expressed in astrocytes (Fig. 5E). This marks the first demonstration that the Cl^−^ conductance is essential for hEAAT1 function *in vivo*. Next, we tested two mutations that are akin to P290R because they have increased Cl^−^ flux (P98G, P392V), but unlike P290R, they have little effect on glutamate transport activity (45, 46). P98G and P392V were expressed well in astrocytes (Fig. 5E), but neither was able to rescue larval crawling (Fig. 5A–D), supporting the idea that increased Cl^−^ flux alone can contribute to hEAAT1 dysfunction. A positive correlation was observed between the effects of the mutations on hEAAT1 Cl^−^ channel function (*E*_*rev*_), and the impact on mutant larval locomotion (Fig. 4D). The larger the change in hEAAT1 Cl^−^ channel function (either a decrease or an increase), the more severe the phenotype observed in the larval crawling assays, suggesting that tight regulation of Cl^−^ flux is critical for EAAT1 function in the intact CNS.

**Figure 5.**
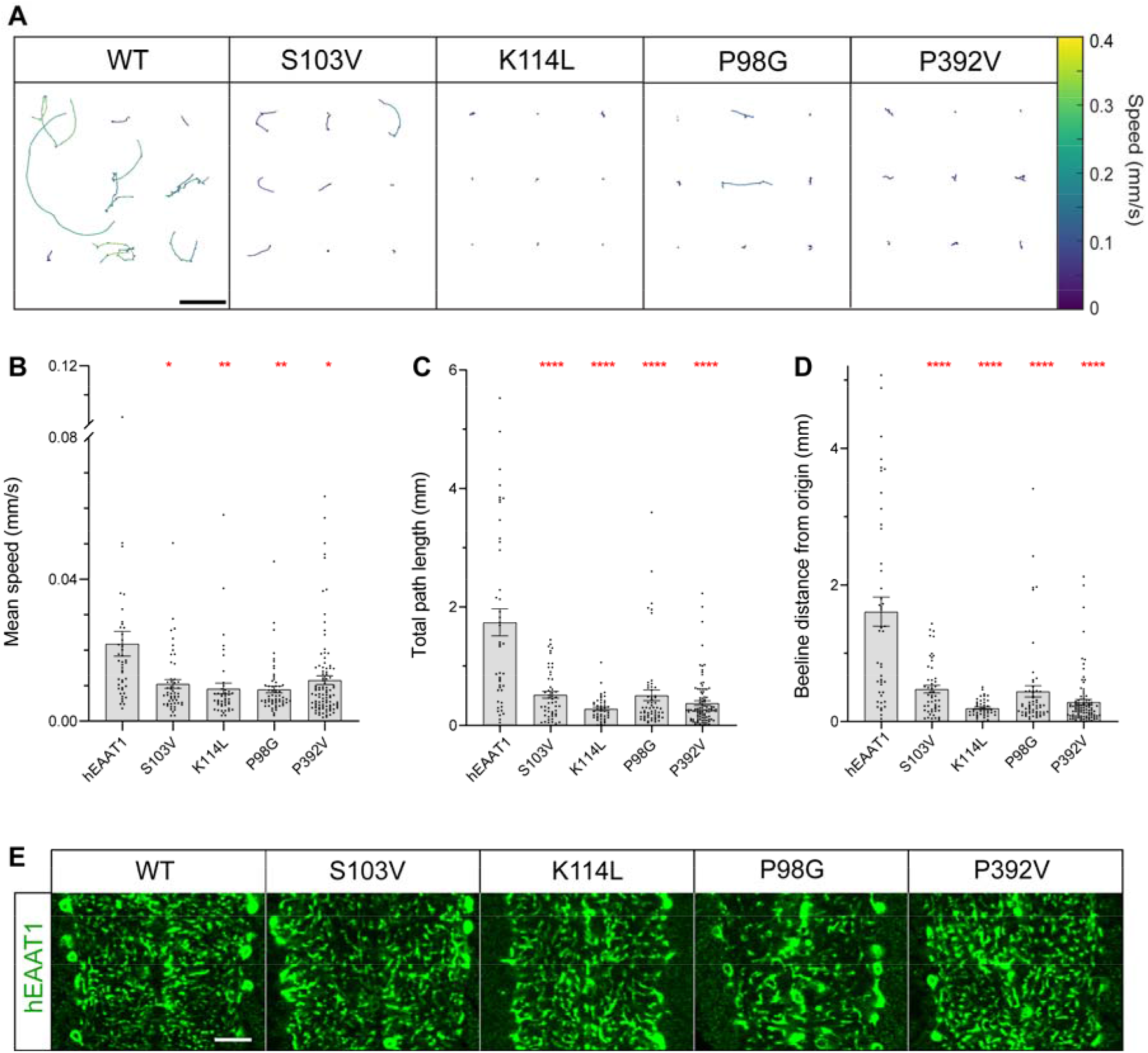
Rescue assay in *Drosophila* demonstrates the hEAAT1 Cl^−^ channel is essential for CNS function. **(A)** Representative trajectories of crawling paths of L1 larvae in 180s, with velocity heatmap. Paths of nine larvae are shown for each genotype where *dEAAT1*-null animals were rescued by hEAAT1-WT or mutations of hEAAT1 known to reduce Cl^−^ channel function (S103V, K114L) or increase it (P98G, P392V). Scale bar (5 mm) in hEAAT1 panel applies to other panels also. **(B-D)** Quantification in bar graphs of mean speed (B), total path length (C), and the beeline distance from origin (D) for larvae over 60s of continuous tracking. One-way ANOVA tests (Brown-Forsythe) were performed for mean speed F(4,106.5) = 7.029, p < 0.0001, for total path length F(4, 74.75) = 26.08, p < 0.0001, and for beeline distance F(4, 73.08) = 27.28, p < 0.0001. **(E)** Immunohistochemistry for hEAAT1 shows that S103V, K114L, P98G and P392V mutations do not appear to affect the expression of hEAAT1 nor its distribution to astrocyte processes within CNS neuropil. Panels show a single optical confocal section within the ventral nerve cord of *dEaat1*-null L1 larvae with astrocyte-specific expression (with *alrm-Gal4*) of hEAAT1 or S103V, K114L, P98G, or P392V. Scale bar =10 μm and applies to all panels.

### Disruption of Na3 binding renders hEAAT1 a non-functional transporter

As no substrate-activated conductance could be measured for M128R, despite adequate expression on the surface of oocytes (Supplementary Fig. S1), we sought to investigate this interesting mutation further. Methionine 128 is located in TMD 3 of hEAAT1, and available structures of SLC1A members reveal that this residue faces the lipid bilayer and is in close proximity to the one of the three Na^+^ binding sites, namely Na^+^ binding site 3 (Na3) (42, 43, 53, 54). The binding of a Na^+^ ion to Na3 is thought to be critical for the substrate transport process (43). For M128R, the substituted arginine side chain is two bonds longer than a methionine side chain, and more importantly, it carries a permanent positive charge. If the side chain of the substituted arginine were to remain in the same position as the methionine, namely, oriented toward the plasma membrane (termed the “OUT” conformation), it would be expected to substantially perturb the local lipid bilayer. Alternatively, the side chain of the substituted arginine might flip towards the protein (termed the “IN” conformation) where the positive charge could disrupt Na^+^ binding at the Na3 site and interfere with the transport process.

To further explore the role of M128, this residue was mutated to a lysine residue (M128K), which is also positively charged but only one bond longer than a methionine side chain. While M128K could transport L-[^3^H]glutamate into oocytes (Supplementary Fig. S3A), it had ~8-fold increase in affinity for l-glutamate (*K*_*m*_ = 3.9 ± 0.3 μM) and ~4-fold increase for l-aspartate (*K*_*m*_=3.0 ± 0.7 μM) and the Na^+^ affinity for M128K (*K*_*m*_ = 33.2 ± 6.2 mM) was ~3-fold lower than hEAAT1 (*K*_*m*_ = 11.0 ± 1.1 mM) (Supplementary Fig. S3B-D and Table 1). Therefore, while the M128K mutation was able to support glutamate transport, the rate of transport was reduced, likely due to weaker Na^+^ binding and increased substrate affinity.

Since a substrate-activated conductance could not be measured for M128R, we investigated Na^+^ interactions by examining the properties of the pre-steady state current of hEAAT1 expressed in oocytes. In the absence of substrate, Na^+^ binding (and unbinding) to hEAAT1 can be indicated by the time (relaxation time, ***t***) the pre-steady state current takes to stabilize to a new equilibrium (steady-state current), after perturbation of the membrane potential (Fig. 6A). A component of this capacitive pre-steady state current has been demonstrated to be from Na^+^ unbinding (and re-binding) and is referred to as the Na^+^ transient current (55).

**Figure 6.**
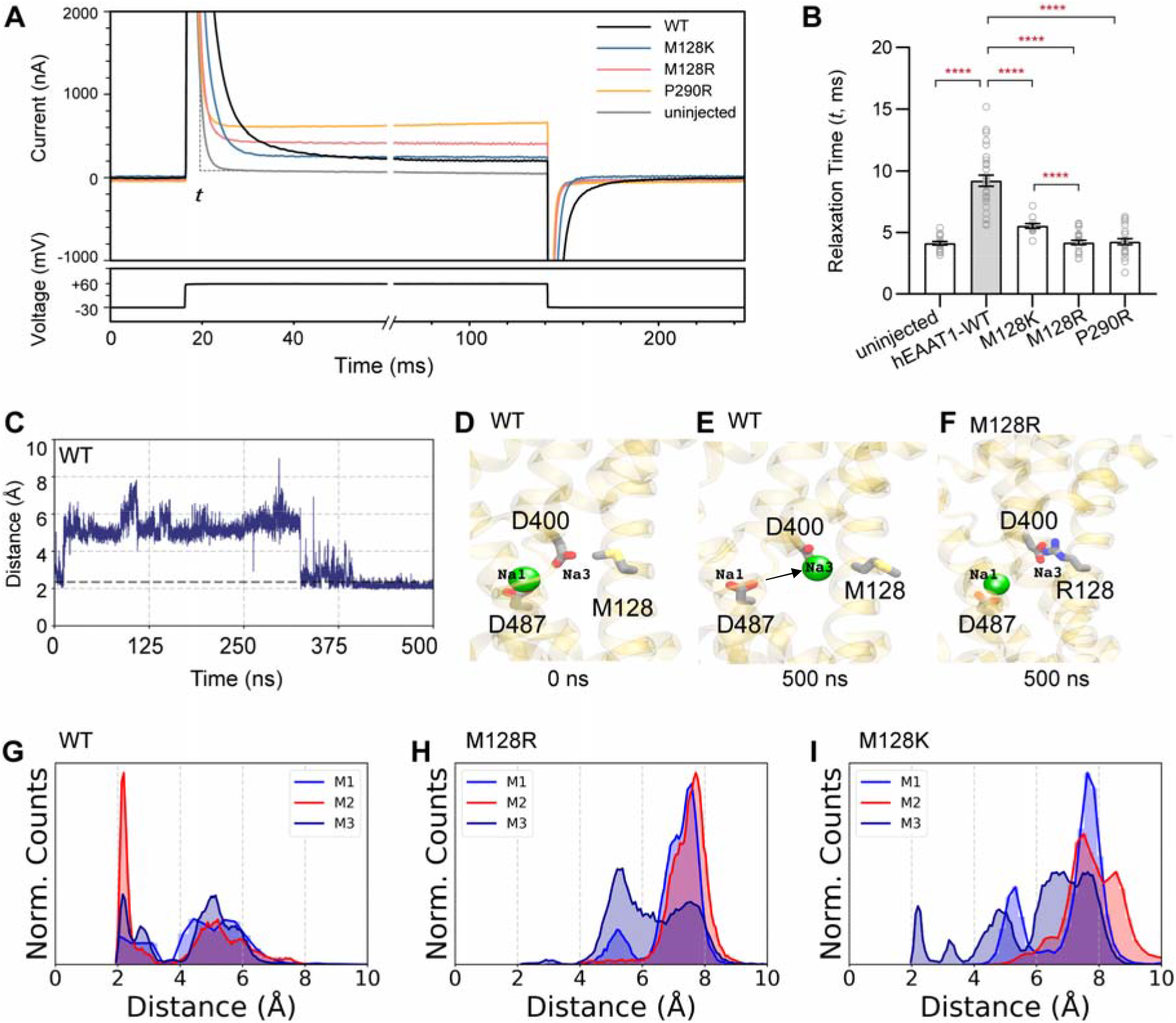
The positive side chain of M128R disrupts Na^+^ transition to site 3 (Na3). (**A**) Representative current trace for hEAAT1, M128K, M128R, P290R and uninjected cells recorded upon voltage jumps (from −30 mV to +60 mV) in 96 mM NaCl buffer are shown. (**B**) The relaxation time (***t***) for the pre-steady state of each recorded current trace was measured. One-way ANOVA tests (Brown-Forsythe) were performed F(4,58.1) = 75.91, p < 0.0001. (**C**) A representative time evolution of one Na^+^-hopping trajectory observed during molecular dynamic simulations of trimeric hEAAT1, in a neuronal membrane. The graph shows the distance between a Na^+^ ion and D400 in the Na3 site. **(D, E)** For hEAAT1-WT, the Na^+^ ion position at Na1 moves towards the Na3 site (indicated by black arrow) where it remained stably bound until the end of the 500 ns simulation. (**F**) For M128R, the Na^+^ ion remains in Na1 during the 500 ns simulation. Major residues involved in Na^+^ coordination in Na1 (D487) and in Na3 (D400) are highlighted in stick representation and only the Na^+^ ion in Na1 is shown for clarity (**G-I**) The distance between the Na^+^ ion and D400 (Na3) was used to monitor movements of the ion from the Na1 to the Na3 site. Distance distribution plots show four independent 500 ns simulations. (**G**) In hEAAT1-WT, we observe a large population with smaller distances (< 3 Å) in all three monomers (M1-M3), suggesting movement of the Na^+^ ion, initially in Na1, to Na3. (**H**) This population is absent in the M128R system, as highlighted by larger distance distributions (> 3 Å). (**I**) In the M128K system, a small population corresponding to the movement of a Na^+^ ion moving from Na1 to Na3 (< 3 Å) is observed.

In oocytes expressing hEAAT1, the Na^+^ transient currents resulting from voltage pulses stepped from a holding potential of −30 mV to +60 mV displayed a slower relaxation of the pre-steady state current (***t*** = 9.2 ± 0.5 ms), compared to uninjected control cells (***t*** = 4.1 ± 0.1 ms) (Fig. 6B), demonstrating an hEAAT1 transporter-specific component of the Na^+^ transient current. Compared to hEAAT1, the average time to relax to the new equilibrium was reduced for transporters with weakened Na^+^ affinity (P290R, ***t*** = 4.3 ± 0.2 ms and M128K, ***t*** = 5.5 ± 0.2 ms), likely due to reduced interaction of Na^+^ with these transporters. Interestingly, M128R reached steady state almost as quickly as control uninjected cells on average (***t*** = 4.2 ± 0.2 ms), demonstrating a loss of the hEAAT1 transporter-specific Na^+^ transient current and supporting the idea that M128R has severely reduced, or no, Na^+^ binding.

To test if the positively charged side chain of M128R affects Na^+^ binding to Na3, all-atom molecular dynamics (MD) simulations were performed on the outward-occluded conformation of hEAAT1 (PDB:5LLU) embedded in a lipid bilayer representing neuronal membranes with substrate (aspartate) and two Na^+^ ions bound, in Na1 and Na2 (Supplementary Fig. S4A, Table 2). For hEAAT1, events were observed where Na^+^ ions “hopped” from Na^+^ binding site 1 (Na1) to Na^+^ binding site 3 (Na3), a step necessary for full occupancy of the transporter at all three Na^+^ sites (Fig. 6 C–E, G Supplementary Fig. S4B, Supplementary Movie 12). For M128R, no such hopping events were observed in any of the simulations, and the Na^+^ ion remained in position Na1 (Fig. 6 F, H Fig. S4C). This may be attributed to the increased positive charge in the vicinity of the Na3 site caused by the substituted arginine at position 128, which adopts an ‘IN’ conformation during the simulations, most likely to avoid unfavorable interaction with the hydrophobic lipid tails. When M128R adopts the ‘IN’ conformation, the positively charged side chain appears to form a strong electrostatic interaction with aspartate 400 (D400), which is important for coordination of Na^+^ in the Na3 site (Fig. S5 B, E). No such interaction was observed for hEAAT1 (Fig. S5 A, D). For M128K, only in one case was a single ion-hopping event observed where a Na^+^ ion moved to the Na3 site, while in other simulation replicas, the Na^+^ ion either remained in its initial (Na1) position or diffused out into bulk solution (Fig. 6I, Fig. S4D). These MD simulations reveal close interactions between D400 and the positively charged side chain of substituted arginine in M128R, and to a lesser extent with M128K, an interaction that can disfavor Na^+^ binding to Na3 and thereby interfere with hEAAT1 function.

### M128R causes membrane deformation linked to a Na^+^ leak conductance

In addition to the coupled substrate transport conductance and the uncoupled substrate-activated Cl^−^ conductance, the EAATs also have a Na^+^-dependent leak conductance which is carried by Cl^−^ ions (56, 57). This Cl^−^ leak current can be observed in oocytes expressing hEAAT1, where there is a sustained steady-state current in the absence of substrate that is not observed for uninjected cells (Fig. 6A). While M218K displayed a leak current similar to hEAAT1, both P290R and M128R displayed leak currents that were larger in amplitude (Fig. 7A). However, unlike hEAAT1 and P290R, the leak current of M128R does not appear to be carried by Cl^−^ based on the following rationale. The *E*_*rev*_ for the P290R leak current shifted to more negative membrane potentials when the anion in the recording buffer was changed from Cl^−^ to the more permeable anion nitrate (NO_3_^−^). This shift was greater than that of hEAAT1 (Fig. 7D), which agrees with the larger steady-state current observed for P290R (Fig. 7A) and suggests the constitutive steady-state currents observed in hEAAT1 and the P290R mutant are carried by Cl^−^ ions. In contrast, the net shift of *E*_*rev*_ for M128R when the anion in the recording buffer was changed from Cl^−^ to NO_3_^−^ was not greater than that for hEAAT1, suggesting the larger tonic leak current observed for M128R was unlikely to be carried solely by Cl^−^.

**Figure 7.**
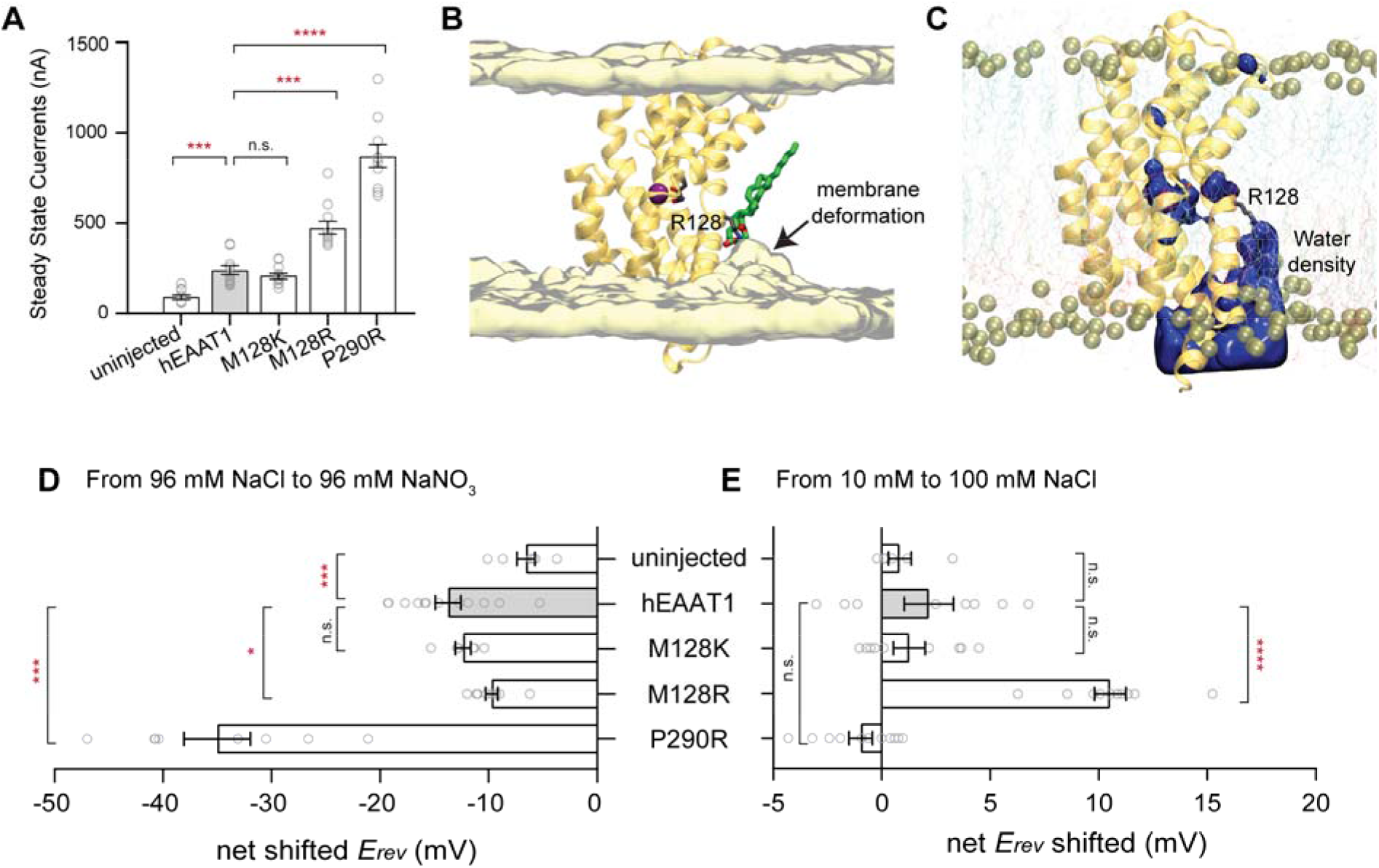
M128R causes membrane distortion. (**A**) The amplitude of the steady-state leak currents of M128R and P290R are larger than hEAAT1 and M128K. For a representative current trace, see **Fig. 6 A**. (**B**) MD simulations revealing membrane deformation when the M128R side chain faces the lipid bilayer, caused by the interactions of M128R (stick representation, grey) with lipid headgroups (an example shown in stick representation, green). Water molecules within 5 Å of the lipids are shown using a yellow surface representation. (**C**) M128R interaction with lipid headgroups results in the recruitment of water molecules into the lipid bilayer. The density of water averaged over last 200 ns of a representative simulation trajectory is shown in blue surface. (**D**) To identify the ions carrying such leak currents, the anion in the recording buffer (Cl^−^) was changed to a more permeable anion (NO_3_^−^), or (**E**) the concentration of Na^+^ was increased by 10-fold. Reverse potentials (*E*_*rev*_) for hEAAT1, P290R, M128R, M128K and uninjected cells were measured, and the changes in *E*_*rev*_ upon alternation in anion (**D**) or cation (**E**) components of the recording buffers were presented as net shift of *E*_*rev*_. One-way ANOVA tests (Brown-Forsythe) were performed for *E*_*rev*_ shifts in anion F(4,12.25) = 48.68, p < 0.0001, and in cation F (4,24.51) = 35.09, p < 0.0001.

During the MD simulations of the M128R mutant, the extended side chain of the substituted arginine was observed to frequently and consistently face the lipid environment in one of the protomers of the hEAAT1 trimer, resulting in considerable deformation of the lipid bilayer in that region (Fig. 7B) accompanied by recruitment of lipid phosphate groups and water molecules into the bilayer space (Fig. 7C, Supplementary Movie 13). This could provide a pathway for ions to leak through the plasma membrane, which may explain the origin of the leak currents observed for M128R and contribute to the pathogenicity of this EA6-related mutation. To determine if the leak current of M128R was carried by Na^+^ ions, sustained steady-state currents were again measured, but under conditions where the concentration of Na^+^ in the recording buffer was changed 10-fold. The net *E*_*rev*_ shift for Na^+^ showed no significant difference between hEAAT1 and uninjected cells (*p* = 0.7369), P290R (*p* = 0.1035) or M128K (*p* = 0.9340). In contrast, the *E*_*rev*_ of M128R shifted by ~10 mV (*p* = 0.0008) to more positive membrane potentials (Fig. 7E), suggesting this tonic steady-state current found in oocytes expressing M128R is carried, at least in part, by Na^+^ ions.

## Discussion

In this study, we characterized EA6-related hEAAT1 mutations using functional studies in *Drosophila* larvae and *Xenopus* oocytes. Our results for A329T and V393I directly linked dysfunction of the hEAAT1 Cl^−^ channel in CNS glial cells with motor deficits *in vivo*. This led us to study other mutations known to increase (P98G, P392V) or decrease (S103V, K114L) hEAAT1 Cl^−^ channel function, which revealed the Cl^−^ channel activity of hEAAT1 is essential for function *in vivo.* Furthermore, our results demonstrate how the EA6-related M128R mutation disrupts Na^+^ binding to hEAAT1, eliminates glutamate transport and Cl^−^ flux, and introduces a leak conductance carried, at least in part, by Na^+^ ions.

The A329T and V393I mutations were identified in multigenerational families with EA in which at least some members had onset of symptoms during childhood. Predictive algorithms for both mutations suggest each is probably pathogenic, though the V393I family had two asymptomatic female carriers of the mutation (36). We found that both A329T and V393I are functional glutamate transporters with only subtle changes in *K*_*m*_ for l-glutamate (A329T), or Na^+^ (V393I). However, both mutations significantly reduced the magnitude of the uncoupled substrate-activated Cl^−^ conductance of hEAAT1, and neither rescued crawling defects in the *Drosophila* model despite robust expression in astrocytes. How these mutations specifically disrupt the hEAAT1 Cl^−^ channel remains speculative but, for V393I, a mechanism can be proposed based on a recent study locating the Cl^−^ permeation pathway in an hEAAT1 homologue, which revealed the Cl^−^ channel is gated by two clusters of hydrophobic residues (31). Removing hydrophobic bulk in these regions through point mutations was found to increase the Cl^−^ conductance, while introducing hydrophobic bulk, decreased the Cl^−^ conductance. Valine 393 is a conserved residue in TM7 and is in proximity to the intracellular hydrophobic gate, therefore substitution for isoleucine at this position (V391I) may introduce an additional hydrophobic barrier for Cl^−^ permeation.

That A329T could partly rescue *Drosophila* crawling is interesting because its effect on the Cl^−^ conductance was milder than that for V393I, correlating the degree to which the hEAAT1 Cl^−^ channel activity is disrupted with the impact of hEAAT1 on motor function. Interestingly, the patient carrying A329T was also found to be heterozygous for an in-frame deletion in *CACNA1A*, which encodes the α1 subunit of the P/Q-type voltage-gated Ca^2+^ channel (12). *CACNA1A* mutations cause episodic ataxia type 2 (EA2) whose clinical features closely resemble those of EA6, and so it is possible that disease in this patient is caused by the combined effects of both mutations.

The C186S mutation was identified in a multi-generational family with EA6 and 1 asymptomatic carrier, while the T318A was found in a patient with no family history of EA6. Curiously, we found no evidence that C186S and T318A mutations are pathogenic, at least on their own, since in our experiments neither one affected hEAAT1 expression, glutamate transport, Cl^−^ channel activity, or larval crawling.

The M128R mutation, likely a de novo mutation, was identified in an EA6 patient whose symptoms first appeared at the age of 11 months (37). M128R was unable to rescue larval crawling and exhibited reduced expression on the surface of *Drosophila* astrocytes (Fig. 2), as reported previously for the P290R mutation (48). When expressed in oocytes, no substrate-activated conductance or L-[^3^H]glutamate uptake could be detected above background, despite sufficient expression at the cell surface. The location of methionine 128 in TM3 is near the Na3 binding site and may inform how a single point mutation can have such drastic effects on transporter function. It has been proposed that binding of a Na^+^ ion to the Na3 site is the first binding event to take place during the transport process, followed by Na1, substrate (glutamate/aspartate), and then Na2 (43). Therefore, potential disruption of Na3 binding by the introduction of a positive charge at M128 may prevent these subsequent binding events. Indeed, our MD simulations revealed that a substituted arginine side chain at position 128 can point toward the lipid bilayer like the native methionine residue or, alternatively, can flip inward to point toward the Na3 site. This prevents hopping of Na^+^ from Na1 to Na3 and thereby blocks subsequent binding events required for the transport process to progress, including entering the Cl^−^ conducting state. Substituting lysine there instead (M128K), which has a slightly shorter side chain but retains the positive charge, did permit glutamate transport, although the apparent affinity for Na^+^ was reduced compared to hEAAT1. In alignment with these functional observations, MD simulations of M128K reveal that Na^+^ hopping events can occur but are less frequent when compared to hEAAT1 likely hindering Na^+^ entry into the Na3 site and the following binding events required for the transport process.

Furthermore, the positive charge of the substituted side chain in M128R may also inform understanding of the ectopic Na^+^ leak conductance observed with this mutation. MD simulations revealed that when the positive side chain of M128R points toward the lipid bilayer, the lipid headgroups are attracted by the positive charge of the arginine reside, distorting the membrane. This appeared to allow water access from the inner leaflet to about half-way across the membrane (near the Na3 site), and we surmise that Na^+^ ions entering the Na3 site can leak through this aqueous cavity when the M128R side chain is pointed toward the lipid bilayer. That an arginine side chain can create a transient cavity within a membrane core allowing water permeation has been reported previously (58), where flipping between distinct orientations of the side chain involved lipid rearrangement and contributed to formation of a pathway for transient water permeation. Such events were not observed with the shorter and less basic lysine side chain (59). Consistent with this, in our MD simulations for M128K the lysine side chain was never observed to point toward the lipid bilayer, perhaps explaining why no Na^+^ leak conductance was observed even though this substitution does impact Na^+^ binding.

Together, our data provide compelling evidence for a link between dysfunction of the hEAAT1 Cl^−^ channel and pathology of the EA6 mutations M128R, P290R, A329T and V393I. Previously, the pathology of the P290R mutation was associated with *gain-of-function* channel activity in hEAAT1, thought to cause excess extrusion of Cl^−^ ions from hEAAT1-expressing glial cells (39, 48, 50). In the patient with the P290R mutation, reduced glutamate recovery may have compounded with increased Cl^−^ flux and contributed to the pediatric onset and severe symptoms. In support of this possibility, it is interesting to note that in our *Drosophila* model, the P290R mutation was shown previously to cause cytopathology in astrocytes and larval paralysis (48), neither of which were observed with the mutations in this study. Although T318A does not seem to be pathogenic on its own, a recent study that co-expressed T318A mutant with wide-type hEAAT1 showed elevated expression levels of the mutant transporter, which consequently contributed to an increase in the Cl^−^ conductance (60). In contrast, both A329T and V393I display significant reductions in Cl^−^ channel activity yet appear to be functional glutamate transporters, while M128R is completely non-functional as a glutamate transporter or Cl^−^ channel but does permit Na^+^ leak through the cell membrane.

These results suggest that a *loss-of-function* of the hEAAT1 Cl^−^ channel activity also contributes to the pathology of EA6, highlighting that too much *or* too little Cl^−^ permeation through hEAAT1 can disrupt astrocytic Cl^−^ homeostasis, cellular membrane potential, and cause CNS dysfunction. This concept is further supported by our findings that mutations that have minimal effects on glutamate transport, but have either elevated (P98G, P392V) or reduced (S103V, K114L) Cl^−^ channel function through hEAAT1, result in *Drosophila* crawling defects like the EA6-mutants M128R, A329T and V393I. The link between the Cl^−^ channel activity of hEAAT1 and the deleterious effects of EA6-related mutations is consistent with the observation that most episodic neurological diseases are channelopathies (61). In EA6 patients with hEAAT1 mutations, altered Cl^−^ channel properties (and Na^+^ leak in M128R) likely disturb ion flux in Bergmann glia. Cl^−^ homeostasis may be restored by other Cl^−^ transport proteins, including hEAAT2, the Na^+^-K^+^-Cl^−^ transporter NKCC (62), the K^+^ and Cl^−^ transporter KCC (63), GABA transporters (GAT) (64), and volume-regulated anion channels (VRAC) (65). Occasions when these systems fail to maintain Cl^−^ homeostasis could explain the episodic nature of EA6 attacks.

## Methods

### Plasmids for expression in Xenopus oocytes

Complementary DNA (cDNA) encoding human excitatory amino acid transporter 1 (hEAAT1) was subcloned into plasmid oocyte transcription vector (pOTV) for expression in *Xenopus laevis* oocytes. Plasmid DNAs were purified with PureLink® Quick Plasmid Miniprep Kit (Invitrogen), linearized with *Spe1* restriction enzyme (New England BioLabs), then transcribed into cRNA by T7 RNA polymerase (mMESSAGE mMACHINE® Kit, Ambion, USA). Mutations identified in EA6 patients were introduced into hEAAT1 plasmid DNA using Q5 site-directed mutagenesis kit (New England Biolabs). Purified plasmid DNAs were sequenced using Big Dye Terminator (BDT) labelling reaction on both strands by the Australian Genome Research Facility (Sydney, Australia). To visualize transporter expression at the plasma membrane of oocytes, DNA encoding enhanced green fluorescent protein (eGFP) was cloned into plasmids encoding wild-type and mutant hEAAT1 transporters, using *BglII* and *EagI* to position eGFP at the C-terminus of hEAAT1.

### Oocyte harvesting and preparation

All chemicals were purchased from Sigma-Aldrich Co. (Sydney, Australia) unless otherwise stated. Female *Xenopus laevis* frogs were purchased and imported from NASCO International (Wisconsin, USA). Oocytes were surgically removed from anaesthetized frogs after administration of tricaine for 12 minutes (buffered with sodium bicarbonate, pH 7.5). Oocytes were defolliculated with collagenase (2 mg/mL) for 1 hour, then stage V oocytes were injected with cRNA (20 ng) encoding the transporter proteins. Injected oocytes were incubated at 18°C in standard frog Ringer’s solution (96 mM NaCl, 2 mM KCl, 1 mM MgCl_2_, 1.8 mM CaCl_2_, 5 mM hemisodium HEPES, pH 7.5) supplemented with 50 μg/mL gentamycin, 50 μg/mL tetracycline, 2.5 mM sodium pyruvate and 0.5 mM theophylline.

### Electrophysiology

2–4 days after injections, currents were recorded using the two-electrode voltage clamp technique with a Geneclamp 500 amplifier (Axon Instruments, Foster City, CA, USA) interfaced with a PowerLab 2/20 chart recorder (ADInstruments, Sydney, Australia) and a Digidata 1322A (Axon Instruments, CA, USA), used in conjunction with Chart software (ADInstuments; Axon instruments). All recordings were made with a bath grounded via a 3 M KCl/ 1% agar bridge linking to a 3 M KCl reservoir containing a Ag/AgCl_2_ ground electrode to minimize offset potentials. Current-voltage relationships for substrate-elicited conductance were determined by measuring substrate-elicited currents during 245 ms voltage pulses between −100 mV and +60 mV at 10 mV steps. Background currents were eliminated by subtracting currents in the absence of substrate from substrate-elicited currents at corresponding membrane potentials. To avoid possible anion loading of the oocytes during repeated application of substrate, experiments determining the *E*_*rev*_ of aspartate-elicited currents were conducted in a buffer with reduced Cl^−^ concentration (10 mM NaCl, 86 mM sodium gluconate, 2 mM potassium gluconate, 1 mM magnesium gluconate, 1.8 mM calcium gluconate, and 5 mM hemi-Na^+^ HEPES) (45). The pH of recording buffers was adjusted using alkaline Tris base to pH 7.5.

The apparent affinity (*K*_*m*_) of glutamate and aspartate for hEAAT1 and mutant transporters were measured by applying increasing concentrations of L-glutamate and L-aspartate in standard frog Ringer’s solution. Substrate-elicited currents (I) at −60mV were fitted to the Michaelis-Menten equation by least squares:

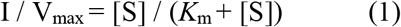

where *K*_m_ is the substrate concentration required to reach half-maximum response, V_max_ is the maximum response, [S] is substrate concentration. In Na^+^ titration experiments, osmolarity was balanced with choline, which does not support transport. L-Glutamate-elicited currents (I) at −60mV in buffers with increasing Na^+^ concentrations were fitted to the Hill equation by least squares:

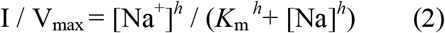

Where *K*_m_ is the Na^+^ concentration required to reach half-maximum response, V_max_ is the maximum response, [Na^+^] is sodium concentration of the recording buffer, *h* is the Hill slope.

### Radiolabelled glutamate uptake assay

Oocytes were incubated in 10 μM l-[^3^H]glutamate for 10 minutes and uptake was terminated by washing oocytes three times in ice-cold standard frog Ringer’s solution. Oocytes were lysed with 1M NaOH and 1%SDS before the addition of 3 mL scintillant (Optiphase HisSafe, PerkinElmer), and l-[^3^H]glutamate transport was measured using a MicroBeta TriLux scintillation counter (PerkinElmer).

### Drosophila stocks and genetics

Standard techniques were used for fly stock maintenance and construction. *Drosophila melanogaster* stocks of *alrm-Gal4*were obtained from the Bloomington Stock Center, and *dEaat1*-null mutants (*Eaat1*^*SM2*^) were published previously (49). To create the *UAS-hEAAT1* transgene, the full-length coding sequence of SLC1A3 isoform 1 (NCBI Reference Sequence: NM_004172.5) was used to order a synthetic gBlock (Integrated DNA Technologies, Inc. (IDT) for hEAAT1 DNA that was codon-optimized for expression in *Drosophila*. The gBlock fragment was inserted into an intermediate vector (pGEX-6P-1) using restriction enzyme digestion and ligation (Not1/Xba1). In the intermediate vector, Q5® Site-Directed Mutagenesis Kit (NEB, Inc.) was used to make point mutations to produce five individual EA6-related amino acid substitutions (M128R, C186S, T318A, A329T, and V393I), plus two substitutions known to reduce Cl*^−^* channel activity (S103V, K114L) and two known to increase it (P98G, P392V) the mutated fragments were then inserted into the UAS expression vector, pJFRC81 (Addgene) (66), with restriction enzyme digestion and ligation (Not1/Xba1). Transgenic flies were generated via standard procedures and φc31-mediated transposition into the *Drosophila* genome at the VK00027 landing site located on chromosome 3 (BestGene Inc.). To select larvae of the correct genotype for immunhistochemistry and behavior assays, fly stocks of *dEaat1*^*SM2*^, *alrm-Gal4(II) / CyO P{Dfd-GMR-nvYFP}; UAS-hEAAT1*^*X*^ / *TM3, P{Dfd-GMR-nvYFP}3,Sb*^*1*^ were intercrossed, where *X* refers to one of the amino acid substitutions noted above.

### Immunohistochemistry

The CNS (brain and ventral nerve cord) of first instar (L1) *Drosophila* larvae of either sex was dissected in Sorensen phosphate buffer (pH 7.2), using standard procedures. Tissue fixation (20 minutes at room temperature) was in formaldehyde (4%) for anti-Gat immunohistochemistry, or in Bouin’s fixative (Polysciences, Inc.) for anti-hEAAT1. Subsequent washes (3X over 30 minutes) were done with Sorensen’s buffer containing 0.1% Triton-X-100 (Sigma Aldrich). Specimens were then blocked and permeabilized with Sorensen’s buffer containing 0.5% Triton-X-100 and 5% normal goat serum (NGS), incubated at 4°C with primary antibodies overnight, washed as above, then incubated with secondary antibodies for 2 hours at room temperature. Both primary and secondary antibodies were diluted in Sorensen’s buffer containing 0.1% Triton-X-100 and 1% NGS. After a final wash as above, specimens were mounted with SlowFade Diamond Antifade Mountant containing DAPI (Invitrogen). Primary polyclonal antibodies used were as follows: rabbit anti-Gat (1:2000; Stork et al., 2014); and rabbit anti-hEAAT1 (1:1000, Synaptic Systems). Secondary antibody used was Alexa Fluor 647-conjugated goat anti-rabbit (1:1000, Life Technologies).

### Microscopy and imaging

*Xenopus* oocytes were examined using Zeiss LSM510 Meta confocal microscope with a 10X objective lens, excited at 488 nm. Averaged fluorescence intensity along the cell surface, which indicates the amount of protein tagged with eGFP on the plasma membrane of each oocyte, were measured using ImageJ and normalized to hEAAT1 level of oocytes from the same batch.

*Drosophila* CNS tissues from *dEaat1*^*SM2*^, *alrm-Gal4(II); UAS-hEAAT1*^*X*^ larvae were examined with an Olympus Fluoview FV1000/BX-63 confocal laser-scanning microscope using a 60X oil immersion objective lens. Captured images were processed using ImageJ software. To observe the localization of mutated hEAAT1 proteins within infiltrative processes of astrocytes, we examined 1μm-thick optical sections of the neuropil, situated in the Z-axis between glial cell bodies flanking the dorsal and ventral surfaces of the neuropil. During analysis, images were thresholded manually to distinguish astrocyte processes from the background. 7-10 larvae were examined in each group.

### Locomotion Behavior

L1 larvae (*dEaat1*^*SM2*^, *alrm-Gal4(II); UAS-hEAAT1*^*X*^) were collected 2-20 hours post-hatching. For each genotype, 5-20 at once were placed into a sieve, rinsed with water, then dried and placed into the center of the arena, a 30 x 30 cm plastic dish containing black ink-stained 2% Bacto agar at room temperature. Movies were captured with a C-MOS camera (FLIR) with a 25 mm fixed focal length lens (Edmund Optics), IR LED strip illumination, and a computer. Hardware modules were controlled through open-source MultiWormTracker (MWT) software (http://sourceforge.net/projects/mwt) (67). Larvae were tracked in real-time at 30 frames/sec for 180 secs in total, including 60s for habituation and 120s of free exploration. Data from multiple runs was pooled for each genotype. Raw videos were never stored. Instead, MWT output text files with the 2D contour (outline) for each larva. These contours were processed with the Choreography module (packaged with MWT) to determine the center of mass for each larva, from which the path trajectory and instantaneous speed (averaged across 21 frames) of each larva was calculated using MATLAB (MathWorks). Custom Java code was used to visualize the crawling behavior described by the MWT data and is available on request. Videos were made from screen-recording these larval contours, processing in Adobe Premiere Pro, then exporting into .mp4 format. The resulting reconstructed videos of 180s were adjusted to 10x real-time speed, giving processed videos of 18s each (Supplementary files). For quantification of locomotion, larvae for which interruptions prevented continuous tracking for 60 seconds within the 120s of free exploration were rejected from further analysis, including animals that crawled out of the field or contacted another larva. Also excluded were those larvae that moved less than one body length. Using the Choreography module and custom MATLAB code that is available upon request, critical parameters of larval crawling motion were calculated over 60s of continuous tracking, including mean speed, the total path length, and the beeline distance from origin. Path trajectories with color heatmaps were generated for representative larvae, where the average instantaneous speed over a moving bin of 0.5 seconds was calculated for each track.

### Membrane Embedding and Equilibrium Molecular Dynamics (MD) Simulations

The simulations system were constructed starting with residues 39-490 of the outward-occluded conformation of human EAAT1cryst-II construct (PDB:5LLU) (44). At first, water molecules were added to the internal cavities of the protein using the Dowser program (68). Protonation states of the titratable residues were assigned using PROPKA (69) and the Membrane Builder module of CHARMM-GUI (70, 71) was employed to embed the trimeric hEAAT1 in a pre-equilibrated patch of neuronal-like membrane containing 1-palmitoyl-2-oleoyl-sn-glycero-3-phosphocholine (POPC), 1-palmitoyl-2-oleoyl-sn-glycero-3-phosphoethanolamine (POPE), 1-palmitoyl-2-oleoyl-sn-glycero-3-phosphoserine (POPS), cholesterol (Chol), 1-palmitoyl-2-oleoyl-sn-glycero-3-phosphoinositol (POPI), and small quantities of PIP1, PIP2, and PIP3 lipids (72). An asymmetric lipid bilayer was used with a lipid ratio of Chol:PC:PE:PS:POPI:PIP2:PIP3 = 45.8:29.2:4.5:17.5:1:1:1 in the cytoplasmic leaflet and a lipid ratio of Chol:PC:PE = 45.6:47.8:6.6 in the extracellular leaflet. The system was solvated and neutralized with 150 mM of NaCl. The final constructed system contained ~256,000 atoms with dimensions of 146 x 146 x 126 Å^3^ prior to any simulations.

All simulations were performed under periodic boundary conditions using NAMD2 (73, 74), CHARMM36m protein and lipid forcefields (75, 76) and TIP3P water model (77). During the initial equilibration phase, protein backbone atoms were harmonically (*k* = 1 kcal/mol/Å^2^) restrained to their initial positions. The restraints were released at the start of the production run. All the non-bonded forces were calculated with a cut-off distance of 12 Å, switching at 10 Å. A Langevin thermostat using γ = 1 ps^-1^ was used to maintain the system temperature at 310 K, and long-range electrostatic forces were calculated using the particle mesh Ewald (PME) method (78). The pressure of the system was maintained at 1 bar along the membrane normal using the Nosé-Hoover Langevin piston method (79). An integration time step of 2 fs was used in all the simulations.

We performed MD simulations on three different protein constructs, namely: hEAAT1, M128R, and M128K. All the mutations were incorporated using the Mutate plugin of VMD (Visual Molecular Dynamics) (80). As we were interested to capture the differential role of WT and mutant constructs on Na^+^ binding events to the Na3 site, we performed four independent 500 ns simulation replicas for each system starting from the partially bound (2 Na^+^ in the Na1 and Na2 sites and the substrate) trimeric state of the protein. All the analysis was performed in VMD using in-house scripts. To monitor the movement of the Na1 ion towards the Na3 site, we measured the distance between the Na^+^ ion and the side-chain oxygen atoms of D400 (negative residue that coordinates Na^+^ in the Na3 site). We also monitored the interactions between the residue at position 128 and D400 by calculating the minimum distance between the terminal atoms of the two side chains, namely, the S atom in methionine, the oxygen atoms in aspartate, and the terminal hydrogen atoms in either lysine or arginine. The water occupancy profiles were calculated using the VolMap Tool plugin of VMD.

### Statistics

GraphPad Prism (v7 or v8) was used to prepare graphs and perform statistical analysis. All values presented as mean ± standard error (SEM) with number of cells used (n) indicated for hEAAT1 kinetic data. Each dot in bar graphs represents the signal response from an individual cell. One-way ANOVA tests (Brown-Forsythe) were performed, with Dunnett T3 post hoc analysis of multiple comparisons, and asterisks in graphs denote the significance of P-values comparing indicated group to controls (*, P<0.05; **, P<0.01; ***, P<0.001, ****, P<0.0001). Bars with no asterisk indicate there was no significant difference from the designated control.

### Study approval

All *Xenopus laevis* surgical procedures have been approved by the University of Sydney Animal Ethics under the Australian Code of Practice for the Care and Use of Animals for Scientific Purposes (protocol 2016/970).

## Author contributions

DJvM and RMR conceived the study, QW, AA, SP, ET, DJvM, RMR designed the experiments, QW, AA, SP, EC, JXZ, performed experiments, acquired data and analyzed data, AG, TO provided and adapted tools, QW, AA, SP, ET, DJvM, RMR wrote the manuscript. All the authors critically reviewed and approved the final version of the manuscript. QW and AA share first authorship, with the order determined by consensus.

## Acknowledgements

This work was supported by the Australian National Health and Medical Research Council Project Grant APP1164494 (RMR, DJvM), the Canadian Institutes of Health Research (DJvM), and the Canada Foundation for Innovation (DJvM), and National Institutes of Health grants P41-GM104601 (ET) and R01-GM123455 (ET). The authors wish to thank Professor R. Vandenberg and Dr. E. Peco for helpful suggestions and sharing reagents, and C. Handford and those that support the *X. laevis* colony at the University of Sydney. Computational resources were provided by XSEDE (grant MCA06N060 to ET), NCSA Blue Waters (ET) and Microsoft Azure (ET).

