## Supplementary Figures for "Ataxia-linked SLC1A3 mutations alter EAAT1 chloride channel activity and glial regulation of CNS function"

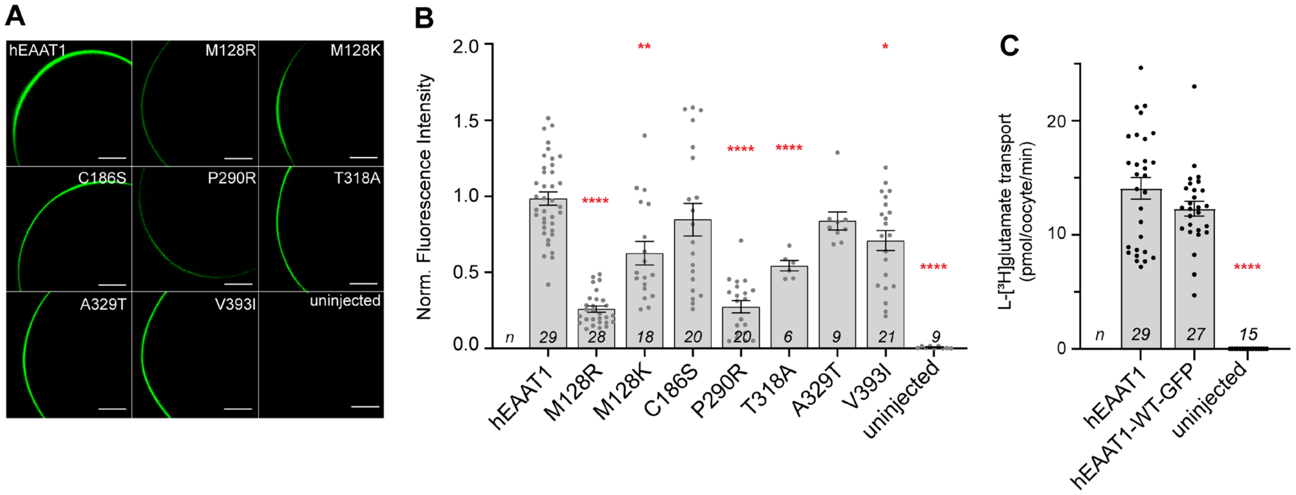


**Supplementary Figure 1. Surface expression of hEAAT1-WT and mutant transporters in *Xenopus laevis* oocytes. (A)** Representative images of oocytes expressing GFP-tagged hEAAT1-WT and mutant transporters are shown. Scale bars indicate 200 µm. **(B)** Average fluorescence intensity was measured along the cell surface to quantify relative expression levels between transporters. **(C)** Rates of L-[$^3$H]glutamate transport were unaffected by addition of GFP tag. Values are normalised to the mean of wild-type values from the same batch of oocytes. Data represented as mean ± SEM, and the number of cells (n) used is indicated. Each dot represents the signal response from an individual cell. Experiments were conducted twice across two different batches of oocytes. One-way ANOVA tests (Brown-Forsythe) were performed for fluorescence intensity $F(8,79.75) = 32.23$, $p < 0.0001$, and L-[$^3$H]glutamate uptake $F(2,48.7) = 94.78$, $p < 0.0001$.

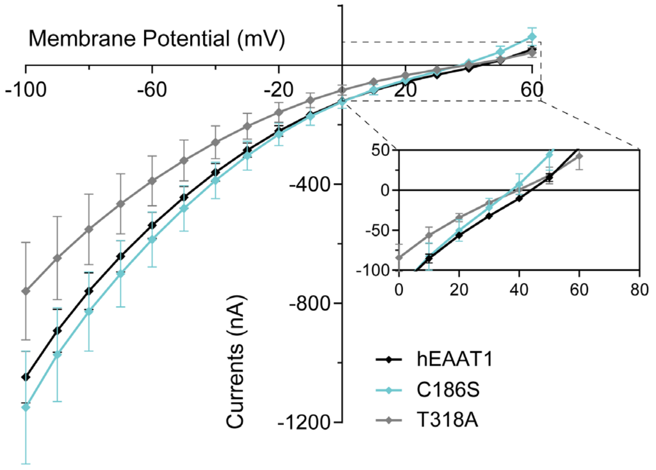


**Supplementary Figure 2. Substrate activated Cl⁻ conductance.** Current-voltage (IV) relationships for hEAAT1, C186S and T318A, where reversal potentials ($E_{rev}$) were analysed in **Fig. 4 B**. Currents were elicited by 100 µM L-aspartate in 10 mM $Cl^-$ buffer, in the presence of 100 mM $Na^+$ (gluconate used as $Cl^-$ substituent to maintain osmolarity).

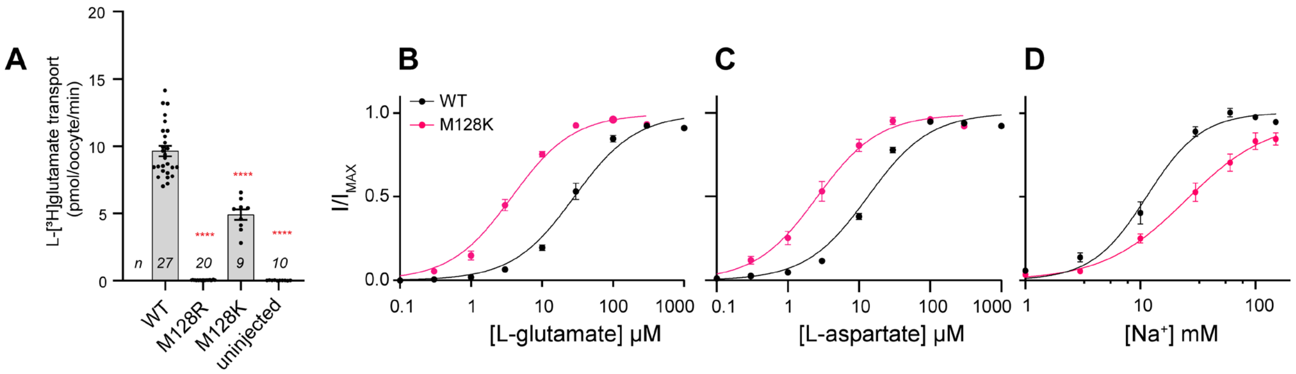


**Supplementary Figure 3. M128K partially restores glutamate transporter function.** (**A**) L-[³H]-glutamate uptake, and dose-response relationships for L-glutamate (**B**), L-aspartate (**C**) and $Na^+$ (**D**) for mutant transporter M128K are shown. $Na^+$ titrations were performed with a saturating dose of L-glutamate (300 µM). Data for hEAAT1, M128R and uninjected cells from **Fig. 3 B-E** is provided for comparison. The number of cells (n) used is indicated in (A) and **Table 1** for panel B-D. One-way ANOVA tests (Brown-Forsythe) were performed for L-[³H]glutamate uptake $F(3,31.8) = 374.7$, $p < 0.0001$.

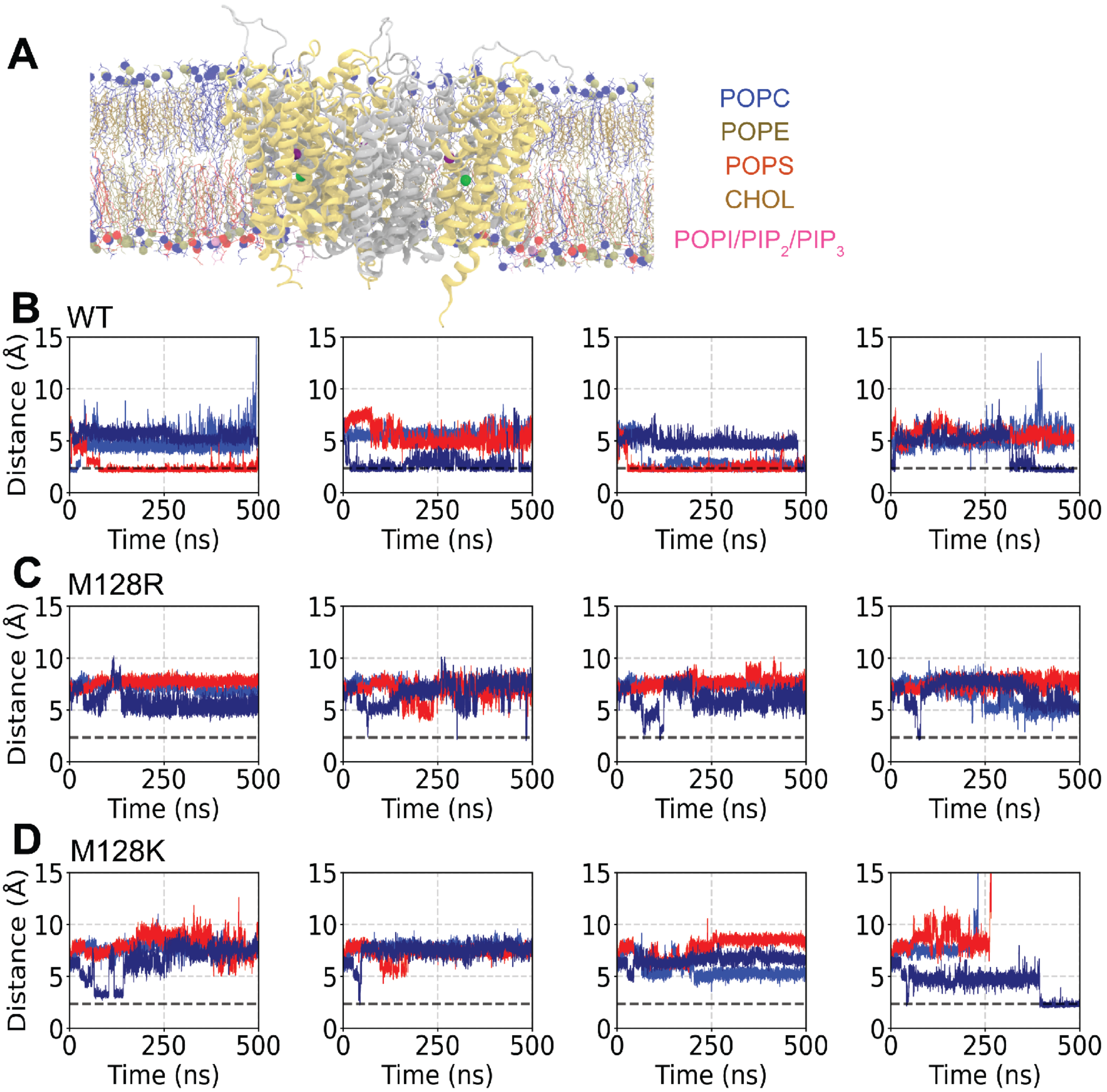


**Supplementary Figure 4. Dynamics of $Na^+$ migration in hEAAT1 and EA6 mutant transporters.** (**A**) Initially the outward-facing partially-bound conformation ($Na^+$ occupying both Na1 (purple spheres) and Na2 (green spheres) sites, with aspartate bound) of hEAAT1, in a trimeric state, was embedded into an asymmetric neuronal lipid bilayer containing POPC (tan), POPE (grey), POPS (red), Cholesterol (cyan), POPI (ice blue), and small quantities of PIP1, PIP2, and PIP3 (grey) lipids. The scaffold and transport domains of the transporter are shown in cartoon representation in silver and gold, respectively. Four independent simulations were run in each system. We monitored the time evolution of $Na^+$ binding to the Na3 site by calculating the distance between the ion initially in Na1 and D400, a negative side chain that coordinates the ion in Na3. (**B**) Multiple $Na^+$ hopping events are captured in hEAAT1, where the $Na^+$ ion from Na1 translocated to Na3 (dashed line). (**C**) In M128R, no movement of $Na^+$ to Na3 is observed. (**D**) In M128K, only one single hopping event was captured.

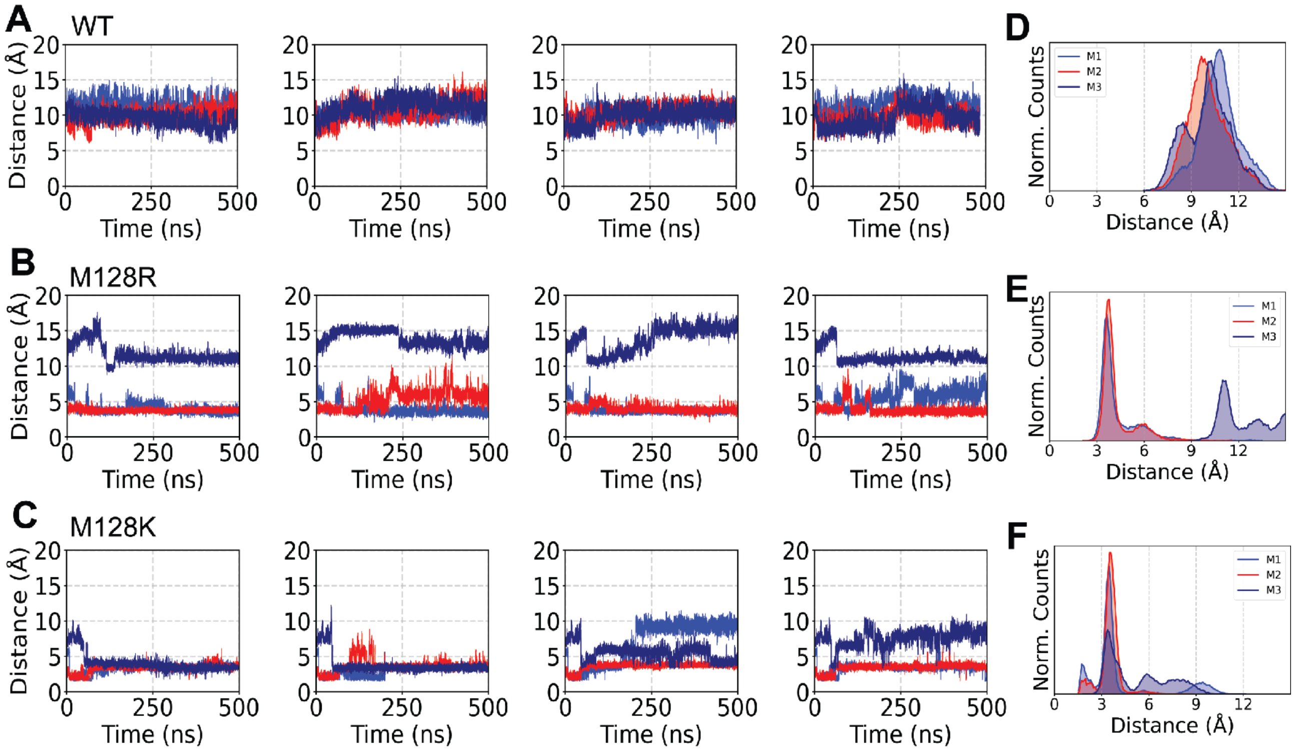


**Supplementary Figure 5. The basic side chains of M128R and M128K can interact with D400.** (**A-C**) Time series of the minimum distance between any side chain oxygen atom of D400 and either the S atom in M128 (**A**), or terminal hydrogens in M128R (**B**) or M128K (**C**) was monitored to represent the proximity of the side chains. Four independent simulations were run in each system, and data for each protomer in each simulation are plotted with a different color. (**D-F**) The distribution histograms of the distances for each protomer (M1-3) in the three systems are shown. (**D**) M128 in all three hEAAT1 protomers adopts an 'OUT' conformation and does not come in proximity to D400, as indicated by all distance distributions longer than 7 Å. (**E**) The arginine side chain in one of the three protomers of M128R also adopts an 'OUT' conformation (M3), while in the other two (M1, M2) it adopts an 'IN' confirmation and forms a salt-bridge with D400. (**F**) The lysine side chain of M128K fluctuates between the two states, with a higher propensity for an 'IN' confirmation. Overall, the presence of a basic side chain at position 128 (M128R or M128K) substantially increases the probability of salt-bridge formation with D400 as illustrated by the distance distributions in **E** and **F**.

**Movies 1–11.** 10x speed reconstructions of infrared tracking of L1 larvae over 180 s, demonstrating lack of larval crawling in dEaat1 null animals (Movie 1), rescue of larval crawling by expression of hEAAT1 in larval astrocytes with alrm-Gal4 (Movie 2), and varying degrees of rescue with each of the EA6-related hEAAT1 mutations M128R, C186S, T318A, A329T and V393I (Movies 3-7) and the additional mutations that affect $Cl^-$ channel function S103V, K114L, P98G and P392V (Movies 8-11)

**Movie 12.** $Na^+$ migration from the Na1 site to the Na3 site in hEAAT1. Starting from the outward-facing partially-bound conformation of EAAT1 (shown in gold, cartoon representation), spontaneous hopping of $Na^+$ (green sphere) from the Na1 site to the Na3 site (residues D487 (Na1), D400 (Na3) and M128 shown in stick representation).

**Movie 13.** Membrane deformation caused by the M128R mutation. The arginine sidechain (stick representation) at position 128 is frequently observed to adopt an orientation facing the lipids, which results in recruitment of lipid headgroups (stick representation with carbon atoms in green) and bilayer deformation (phosphorous atoms are shown as light brown spheres).
